# μSPIM: A Software Platform for Selective Plane Illumination Microscopy

**DOI:** 10.1101/2020.06.15.151993

**Authors:** Daniel Saska, Paul Pichler, Chen Qian, Chrysia Pegasiou, Christopher L. Buckley, Leon Lagnado

## Abstract

Selective Plane Illumination Microscopy (SPIM) is a fluorescence imaging technique that allows volumetric imaging at high spatio-temporal resolution to monitor neural activity in live organisms such as larval zebrafish. A major challenge in the construction of a custom SPIM microscope is the control and synchronization of the various hardware components. Here we present a control toolset, μSPIM, built around the open-source MicroManager platform that has already been widely adopted for the control of microscopy hardware. Installation of μSPIM is relatively straightforward, involving a single C++ executable and a Java-based extension to Micro-Manager. Imaging protocols are defined through the μSPIM extension to Micro-Manager. The extension then synchronizes the camera shutter with the galvanometer mirrors to create a light-sheet that is scanned in the z-dimension, in synchrony with the imaging objective, to produce volumetric recordings. A key advantage of μSPIM is that a series of calibration procedures optimizes acquisition for a given set-up making it relatively independent of the optical design of the microscope, or the hardware used to build it. Two laser illumination arms can be used while also allowing for the introduction of illumination masks. μSPIM allows imaging of calcium activity throughout the brain of larval zebrafish at rates of 100 planes per second with single cell resolution as well as slower imaging to reconstruct cell populations, for example, in the cleared brains of mice.

## I. Introduction

Selective/Single Plane Illumination Microscopy (SPIM) is a powerful method for 4D imaging of biological samples at a high spatio-temporal resolution (Ahrens et al., 2013; Power and Huisken, 2017). A sheet of light a few microns thick is used to excite fluorescent molecules reporting the structure and/or function of the tissue and the images are acquired by a high-resolution camera focused on that same plane. Illuminating a single thin plane minimizes out-of-focus fluorescence to allow the visualization of individual cells and their processes at illumination intensities that minimize photobleaching and phototoxicity. Rapid movement of the sheet through the tissue then allows for collection of complete volumes at high rates. The ability to image with single-cell resolution at depth has made SPIM particularly useful for the study of developmental processes in embryos of C. *elegans* (Rieckher et al., 2015), Drosophila (Keller et al., 2010) and zebrafish (Kobitski et al., 2015; Wan et al., 2019), as well as imaging neural activity in Drosophila embryos and larvae (Chhetri et al., 2015; Lemon et al., 2015), C. *elegans* (Ardiel et al., 2017) and zebrafish (Ahrens et al., 2012; Ahrens et al., 2013; Weisenburger and Vaziri, 2018). This form of light-sheet microscopy also allows for faster imaging in fixed tissue that has been “cleared” to homogenize the refractive index and reduce scattering. Imaging through the cleared brains of mice, for instance, enables the visualization of specific neuronal circuits (Chung and Deisseroth, 2013; Mano et al., 2018).

A number of different implementations of SPIM have been developed (Keller and Ahrens, 2015), but they most commonly use an illumination arm to create a 2D plane of illumination and an orthogonal collection arm that is forming the image onto the camera. There are two main variations on this basic design: the method of light sheet formation and its movement relative to the sample. The original approach creates a stationary light sheet using a cylindrical lens and then translates or rotates the sample with a moving stage. This implementation has been particularly useful in studies of development (Huisken and Stainier, 2009), but it is too slow to monitor the activity of neurons using, for instance, genetically-encoded calcium indicators. Applications in functional neuroscience therefore favour a configuration in which the sample is kept stationary while the light sheet is created using a fast scanning mirror that moves the light beam across a plane at least once per imaging frame, with a secondary mirror moving the beam in the z-dimension (**Figure 2B**). This method has allowed “brain-wide” imaging of neural activity in live zebrafish with single neuron resolution and acquisition frequencies of ∼1 Hz (Ahrens et al., 2012; Ahrens et al., 2013).

Several designs of SPIM have been published but probably the most accessible in terms of both hardware and software are the OpenSPIM (Pitrone et al., 2013) and Open SPIM microscopy (Gualda et al., 2013) projects. Both these implementations are focused on imaging of developmental processes using the slower “moving sample” configuration and therefore have limited use if the aim is to image neural activity through volumes of the brain. The “moving light-sheet” configuration is more complex to control because it requires synchronization of several hardware components with millisecond temporal accuracy. For instance, movements of the imaging objective in the z-dimension must be synchronized with movement of the light-sheet to stay focused on the plane of illumination, often involving 50-100 planes per second. Perhaps for this reason, open software tools for the control of SPIMs in functional imaging experiments are not easily available. We aim to fill this gap by providing a control toolset, μSPIM, which adopts an open software approach for control of a SPIM microscope in which the light-sheet is scanned through a stationary sample. We achieve this by building μSPIM around Micro-Manager (Edelstein et al., 2010), an open-source platform widely used for control of microscopes, to provide a comprehensive user interface (**Fig 3**) with a smooth learning curve. μSPIM has been designed to be used to control a range of custom-built microscopes, for which it is calibrated using semi-automated procedures. We demonstrate the utility of μSPIM for whole-brain imaging in larval zebrafish and neuron reconstruction in cleared mouse brains.

## II. Design Requirements

We start by listing the basic objectives of software controlling a SPIM microscope in relation to the microscope we have constructed (shown in **Figure 1**, see Methods). 1. The illumination arm requires the control of the laser light-source, using either an internal and/or external shutter. Dual-colour imaging requires two independent laser sources to be combined. Depending on the laser model, power may be controlled on long time-scales using USB or RS232 interfaces and modulated on short time-scales by analogue signals, while shuttering requires digital signals. 2. The beam has to be scanned in the x and z dimensions using fast galvanometer mirrors driven through analogue inputs. 3. The collection arm must follow the beam and collect images that are in focus at different z-positions, which is achieved by using a piezo-electric mount for the objective, again controlled through an analogue signal. Finally, 4. The camera acquisition must be accurately timed to collect frames at each z position, which in turn requires accurate calibration of the z mirror so that the z position of the light-sheet is set from the command signal. The software should then allow control of several features of the camera, including frame integration time, pixel binning and gain. Furthermore it should support saving and retrieving images and movies, zooming, defining ROIs, quick and easy changes in imaging parameters and the ability to define various imaging protocols using different sequences of laser illumination

**Fig. 1:**
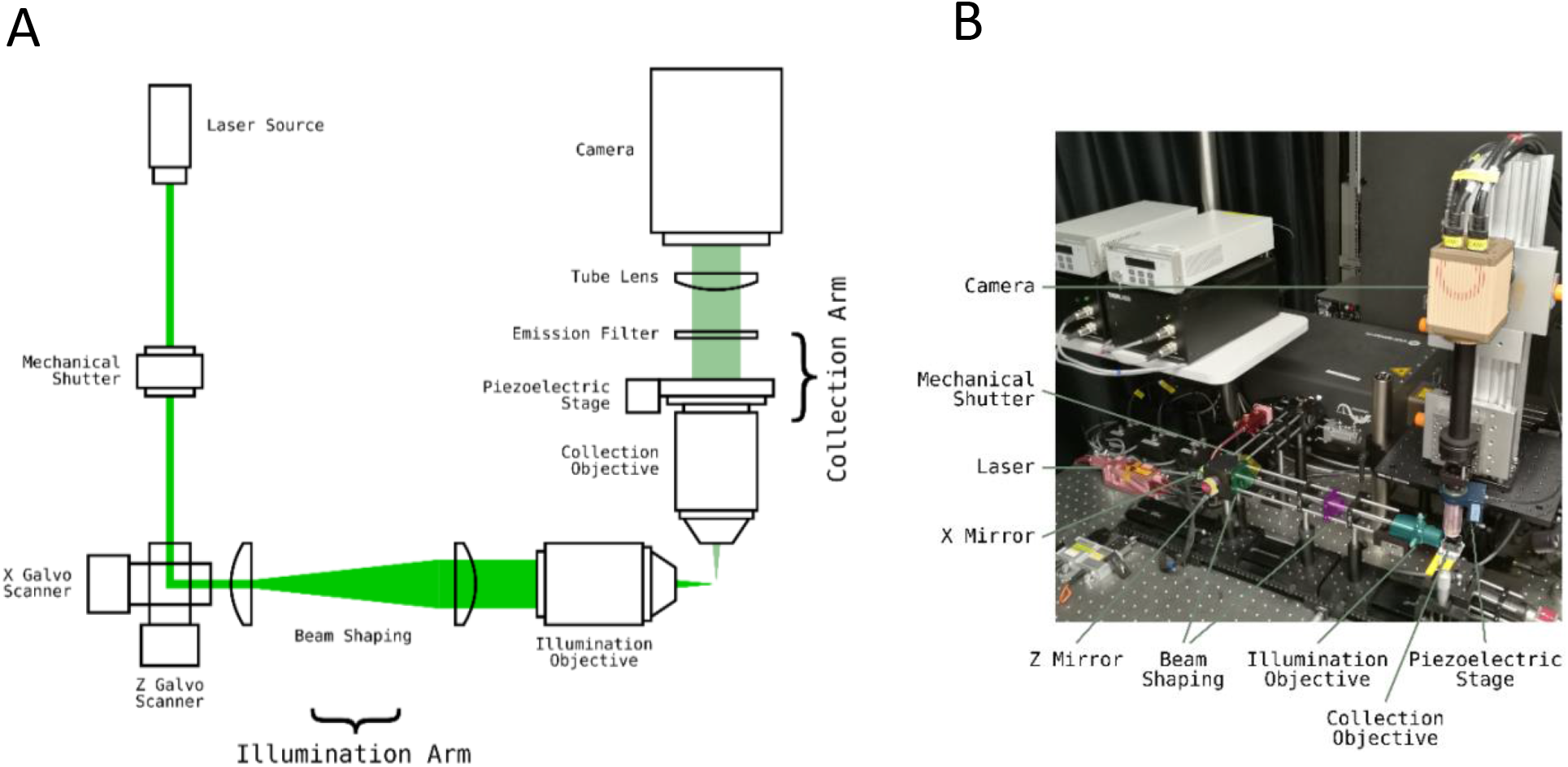
Light-Sheet Microscope Implementation. **(A)** Diagram outlining light-sheet microscope with two mirror galvanometers and a stationary sample. The light sheet is created from a laser beam by the ‘X Galvo mirror’. The ‘Z Galvo mirror’ then moves the sheet through the sample to create volumetric excitation. This is synchronized with a Piezoelectric stage that moves the imaging objective so that the excitation plane always coincides with the imaging plane. **(B)** A photo of an example light-sheet microscope setup with one light-sheet path. The main components are colour coded.

We built μSPIM around the open-source software Micro-Manager (Edelstein et al., 2010), which is based on ImageJ (Schindelin et al., 2015), because it immediately offers the ability to integrate different hardware from a wide range of manufacturers through specific plugins, including a range of cameras, light-sources and shutters required for a SPIM. It provides an integrated environment for image acquisition and a very wide range of post-acquisition processing capabilities through ImageJ. Micro-Manager does not, however, provide hardware triggering, analogue control and monitoring with the precision required for volumetric imaging of neural activity. For this purpose, we used a National Instruments DAC card (NI PCIe-6738) and wrote an executable ‘*μSPIM Control*’ to allow interaction between Micromanager and hardware through its own user interface. μSPIM Control synchronizes internal laser shutters, x and z mirrors, the objective piezo stage and all camera triggering, leaving laser power, external mechanical shutters and camera operating properties under control of Micro-Manager, see Figure 2A. μSPIM therefore provides both accessibility and versatility, retaining the user’s ability to select hardware best suited for the particular setting, as long as it supported through a Micro-Manager plug-in (The current range can be viewed at https://micro-manager.org/wiki/Device%20Support).

**Fig. 2:**
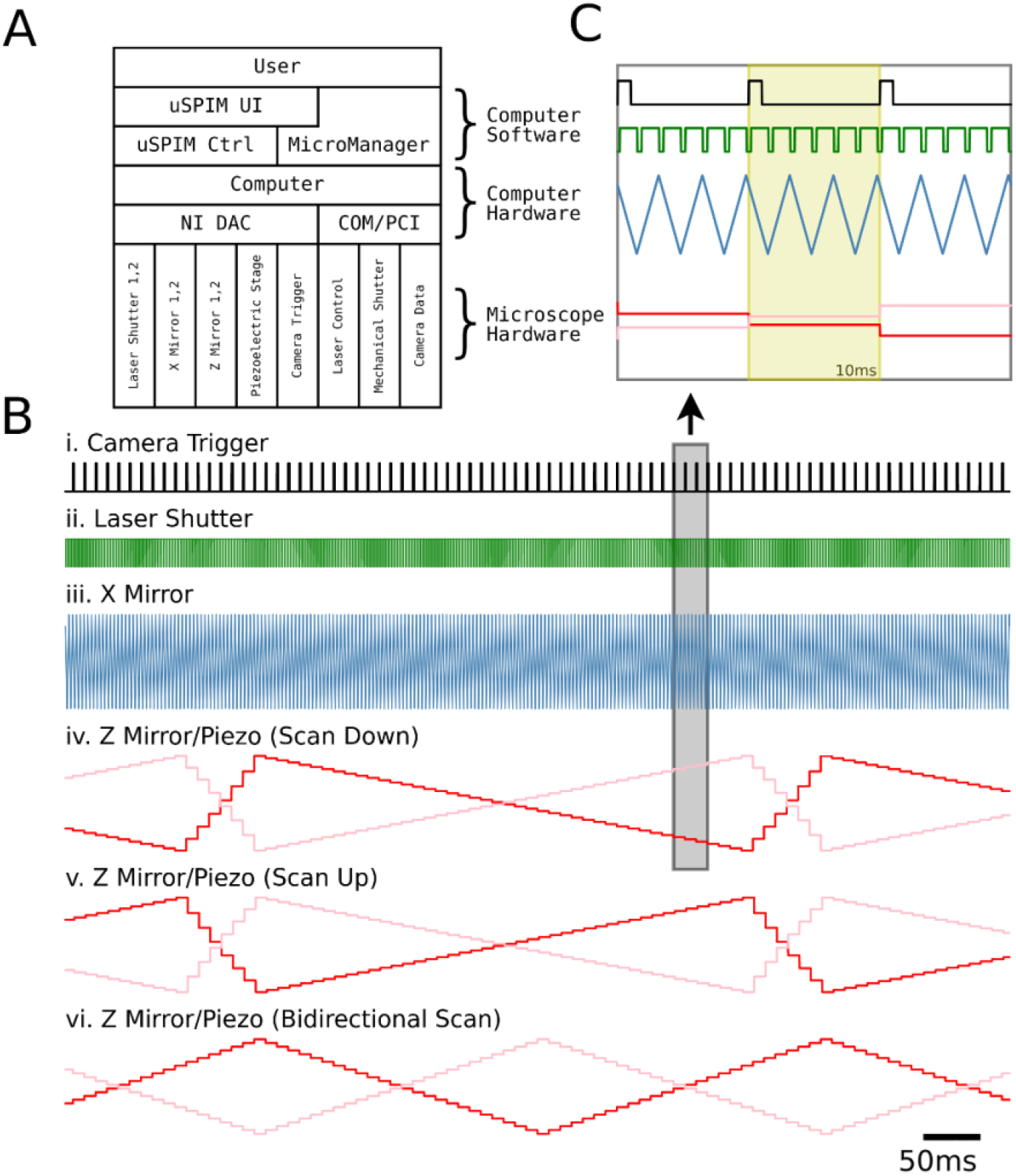
Light-Sheet Microscope Hardware Control: **(A)** Interaction of different components of the μSPIM setup in a typical acquisition setup. μSPIM provides control and synchronization of hardware through NI DAC with MicroManager controlling the camera and mechanical shutters through PCI and COM ports. **(B)** Traces of the command signals for a volume acquisition with 43 planes showing camera trigger (i.), laser shutter (ii.), X mirror signal (iii.), Z mirror (red) and Piezo (pink) signals for Scan Down (iv.), Scan Up (v.) and Bidirectional (vi.) acquisition modes generate by the μSPIM control software. **(C)** Magnification of boxed region in B with yellow region showing a single plane signal. Laser shutter signal is a result of a recording with Edge Masks enabled.

## III. Principles of operation

The μSPIM toolset consists of two main components: a Java plugin for MicroManager which facilitates all user interaction and configuration, and a C++ control executable which interacts with the National Instruments board to produce hardware control signals **(Figure 2)**. These support the control of two scanning X mirrors for light sheet formation, two scanning Z mirrors and one piezoelectric stage for a Z motion of the collection objective, two laser shutters for laser masking and one trigger for the camera acquisition control and synchronization with other hardware. After an appropriate setup, the Java plugin facilitates all necessary tools for calibration and control of the hardware, providing the users with an intuitive interface without requiring the need for specialized computer knowledge or programming, while allowing flexibility provided by MicroManager. **Figure 2** illustrates the control diagram, outlining the interaction between the user and the light-sheet microscope hardware (**Figure 2A**) as well as the temporal sequence of the control signals (**Figure 2B and C**).

A typical user interface for data acquisition is presented in **Figure 3**, with the μSPIM Control window at the top (A) and MicroManager windows below (B-D). The primary Micromanager window (C) provides access to all functionality related to the acquisition, such as live mode, binning, ROI selection or exposure time. The μSPIM plugin (A) provides access to the variables used for light-sheet formation, such as light-sheet width, volume depth or number of planes as well as providing a set of calibration procedures described in the Calibration section below. During scan mode, the image can be viewed in the classical MicroManger *live* window (D) which also allows the acquisition of snapshots. While acquisition can be manually started through the MicroManager interface, the μSPIM plugin interface provides an acquisition routine executing a synchronized start of acquisition and illumination, making the process of acquiring data (namely volumes) simpler and more robust.

**Fig. 3:**
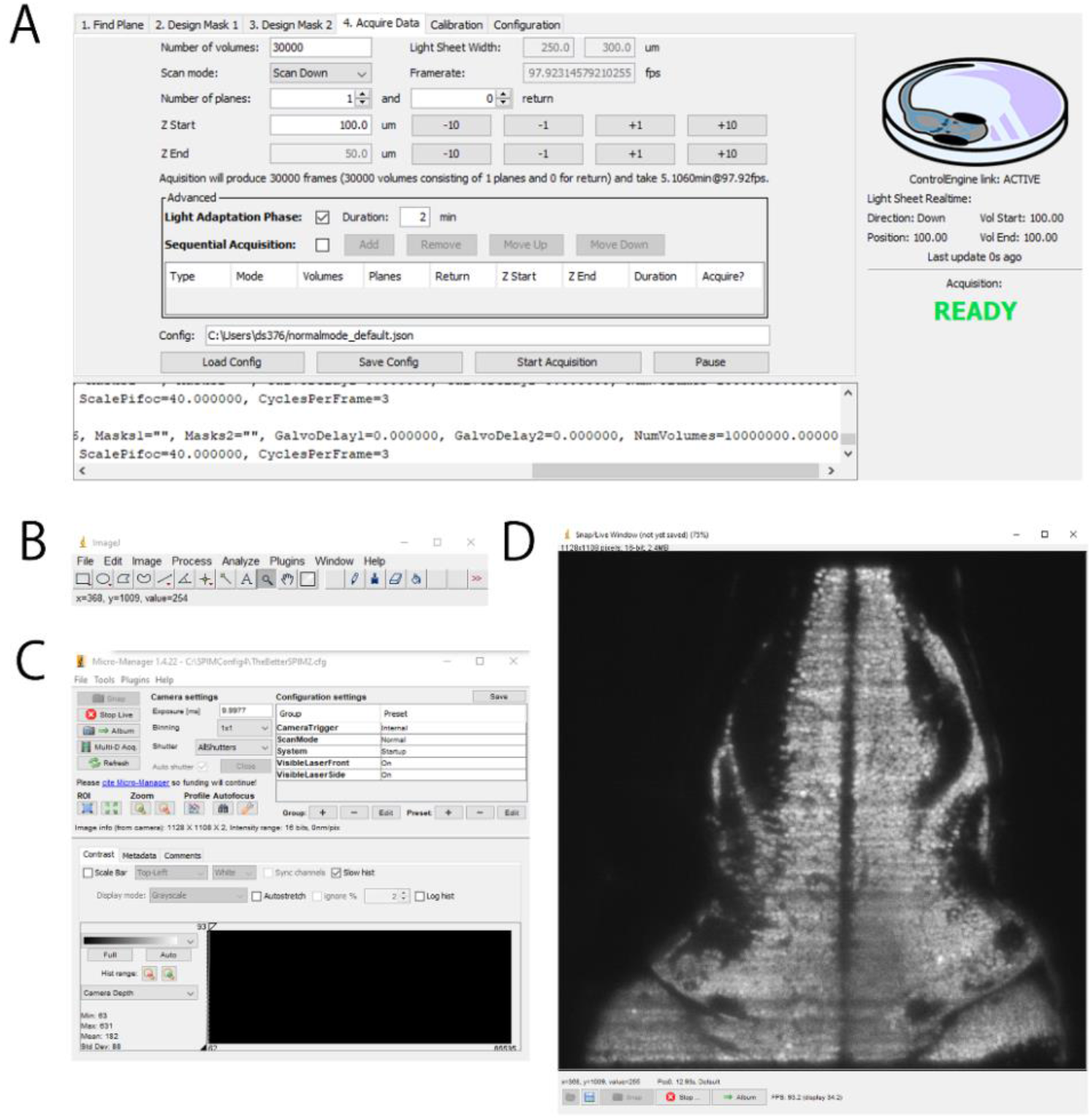
μSPIM User Interface. Following a common MicroManager design, the control interface is separated into several components: μSPIM-provided plugin window with control over the light-sheet generation including calibration and acquisition routines shown in A, basic ImageJ tools shown in B and MicroManager interface providing control over the acquisition hardware shown in C and live view of the camera shown in D.

Taking full advantage of the stationary sample and fast acquisition speeds, we used μSPIM to monitor the activity of neurons across most of the brain of larval zebrafish expressing the nuclear localised Ca_2+_ reporter GCaMP6f panneuronally (Tg(Huc:H2B - GCaMP6f), **Fig. 4**). Larvae were embedded so that the laser excited the sample from the side (arrows in **Fig. 4A**). The width of the sheet spanned 500 μm and covered the entire hindbrain, the cerebellum and parts of the optic tectum. Single planes were recorded at a frequency of 98 Hz. The volume was set to cover a distance of z = 100 μm and consisted of 50 planes with a spacing of 2 μm. The piezo returned to its original position within 10 frames yielding an overall imaging frequency of 1.6 volumes per second. **Figure 4C** shows three representative sections at depths of 20, 30 and 40 μm, respectively. Individual cell bodies are clearly visible in the cerebellum, the anterior and the posterior hindbrain, respectively (**Fig 4D**) and their activity over time is depicted in **Fig 4E**.

**Fig. 4:**
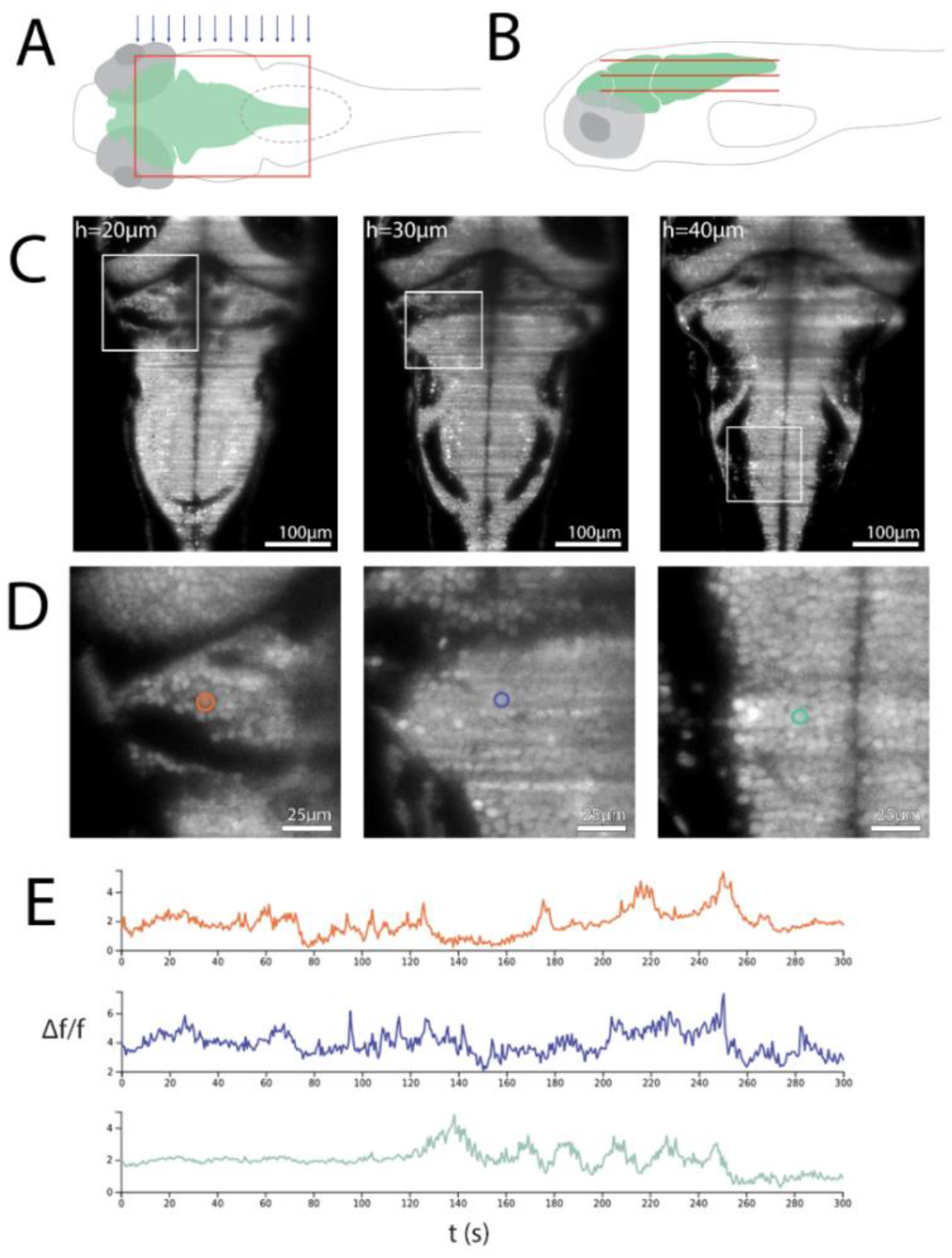
Imaging larval zebrafish: **(A and B).** Schematic of a single light sheet covering the hindbrain of a larval zebrafish from above (A) as well as a number of light sheets (constituting a volume) from the side (B). Sheet size and spacing are not to scale. **(C)** Transgenic zebrafish expressing the calcium reporter GCaMP6f in the cell bodies of all neurons (Tg(HuC:H2B - GCaMP6f)) were embedded in agarose and positioned so that the laser entered the brain from the side (blue arrows in A). Representative sections of the volume taken at 20, 30 and 40 μm depths at a frequency of 1.6 volumes/sec and an integration time of 10.2 ms per section. The step size was 2 μm, hence representative sections are 5 sections apart. The laser was set to 1.8 mW. Scale bar indicates 100 μm. **(D)** magnified view of boxed areas in C. Single cell bodies are clearly visible and three examples are highlighted. **(E)** Activity of single cells highlighted in D.

Many problems in neuroscience require the mapping of neurons and their connections through large volumes of the brain and SPIM has become very useful for such applications because it allows high-speed volumetric imaging of fixed and stained tissues with two key advantages over techniques such as confocal microscopy and multiphoton microscopy - faster speed and much-reduced photobleaching (Stefaniuk et al., 2016; Tomer et al., 2014). Speed improvements of more than 100-fold derive from capturing the whole image simultaneously rather than by scanning a point source, while photobleaching is minimized by only illuminating the plane of interest. This application of SPIM has recently become more important with the development of tissue preparation techniques such as iDISCO - immunolabeling-enabled three-dimensional imaging of solvent-cleared organs (Renier et al., 2014). These techniques remove variations in refractive index that would normally degrade images while allowing visualization of expressed fluorescent proteins or antibody-labelling throughout intact tissues and come into their own when used with SPIM (Dodt et al., 2007; Tomer et al., 2014). μSPIM is also ideally suited to this application and example images are shown in **Fig. 5**, were obtained from the cleared brain of a mouse prepared by the iDISCO technique. Here we used *μSPIM* to control two lasers for dual-colour imaging in red and green channels. The images show excitatory neurons (pyramidal cells) in the primary visual cortex V1 expressing the green calcium reporter GCaMP6f and/or phosphorylated tau protein. Individual neurons are easily visualized and their processes can be followed. Images at this resolution, using planes just 1 μm apart, could be imaged through the brain at speeds of 20 μm/s.

**Fig. 5:**
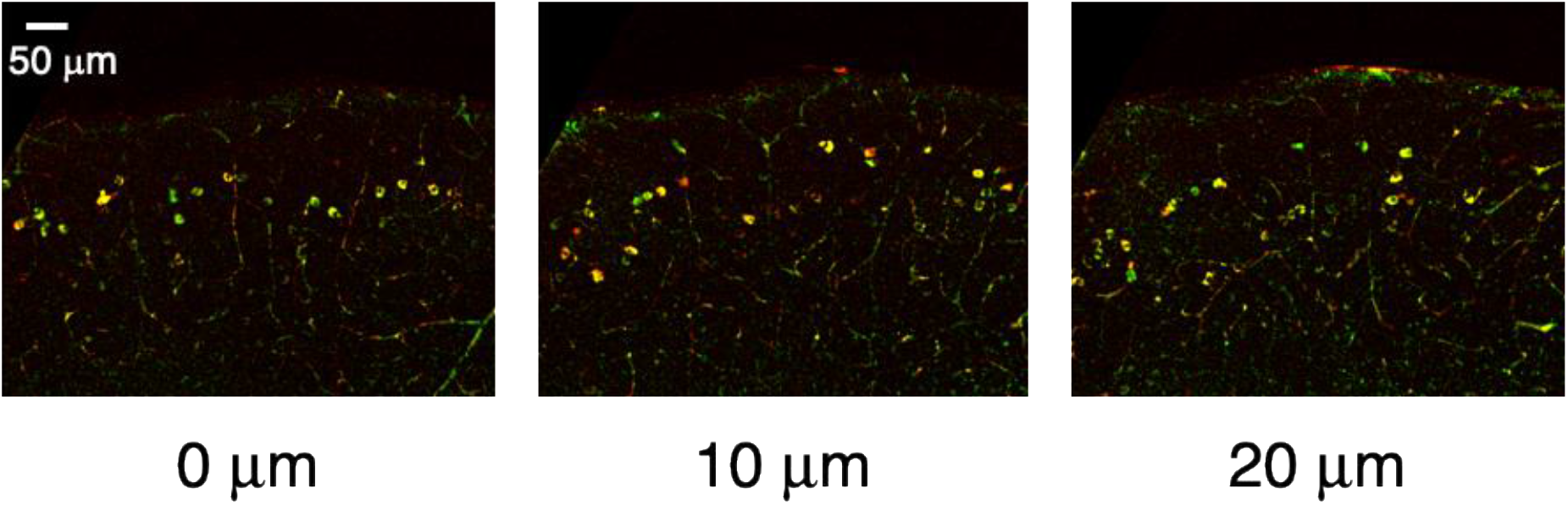
Dual-colour imaging through a cleared mouse brain. The images show excitatory neurons (pyramidal cells) in the primary visual cortex V1, either expressing the GFP-based calcium reporter GCaMP6f (green) and/or tau fibrils (red, labelled with the AT-8 antibody). The sequence of images is spaced 10 μm apart and each is an average intensity projection of 2 planes 1 μm apart. Individual frames were acquired with an integration time of 50 ms. Cell bodies and thin processes of pyramidal neurons can be seen. Cells expressing both GCaMP6f and tau fibrils appear yellow. The surface of the cortex is at the top.

## IV. Computing Requirements

The μSPIM relies only on minimal third-party software for its operation. This includes MicroManager for integration of acquisition hardware (laser control, light path shutter and camera) and National Instruments drivers, in order to interface with the National instruments DAC card. Naturally, drivers and accompanying software must be installed for the selected hardware (such as lasers or camera) in order to ensure MicroManager can communicate with those parts properly. Due to the continuous high data throughput, it is recommended to use a secondary computer for stimulation and/or other intensive tasks to avoid potential performance issues.

The computer specifications necessary for optimal function and acquisition of the setup are highly dependent on the acquisition requirements. For large field of view and high acquisition speeds, the acquisition produces large datasets, making the process highly storage-dependent. State-of-the-art acquisition cameras are able to reach 4 megapixel resolution at 100 Hz and 16 bit depth, producing a theoretical throughput of 838.9 MB/s. This poses a write speed requirement higher than that which can be satisfied by a regular SATA3 storage interface which is limited to a theoretical maximum of 600 MB/s. It is therefore highly recommended to utilize a faster storage solution, such as fast PCI-based solid state disk (SSD) storage (e.g. Intel Optane 905p) or hardware SSD RAID. The acquisition is not highly CPU dependent and neither μSPIM nor MicroManager take a significant advantage in parallel processing. A state-of-the-art consumer line 4- or 6-core CPU is sufficient for optimal performance.

μSPIM supports control of two independently calibrated laser paths. Each path consists of a visible light laser, two scanning mirrors and one laser path shutter (**see Fig. 1**). The laser path shutter must be supported by MicroManager and is used to block light between periods of acquisition and thus reduce photo-bleaching of the sample. One of the two scanning mirrors is used for light sheet formation and the second is used for Z movement. The laser should provide digital or analogue shuttering capability. All other devices are controlled using an analogue signal produced by the National Instrument DAC adapter and should support inputs in appropriate voltage ranges (commonly ±10 V). μSPIM allows for incomplete configurations e.g. single light path with unmodulated laser and single sheet-generating scanning mirror would provide minimal functionality, however most of the features of the μSPIM toolkit would be unavailable. To support the full functionality, the National Instruments DAC card must support at least 8 analogue outputs with no need for analogue or digital inputs as the control is fully feed-forward and does not require any feedback from the hardware.

## V. Calibration

The spatial scales of the signals are greatly dependent on the chosen hardware and thus require the user to perform set of calibration procedures to ensure appropriate alignment of the hardware. Here and in **Figure 6** we outline the principles of the calibration of the different components affecting the acquisition.

**Fig. 6:**
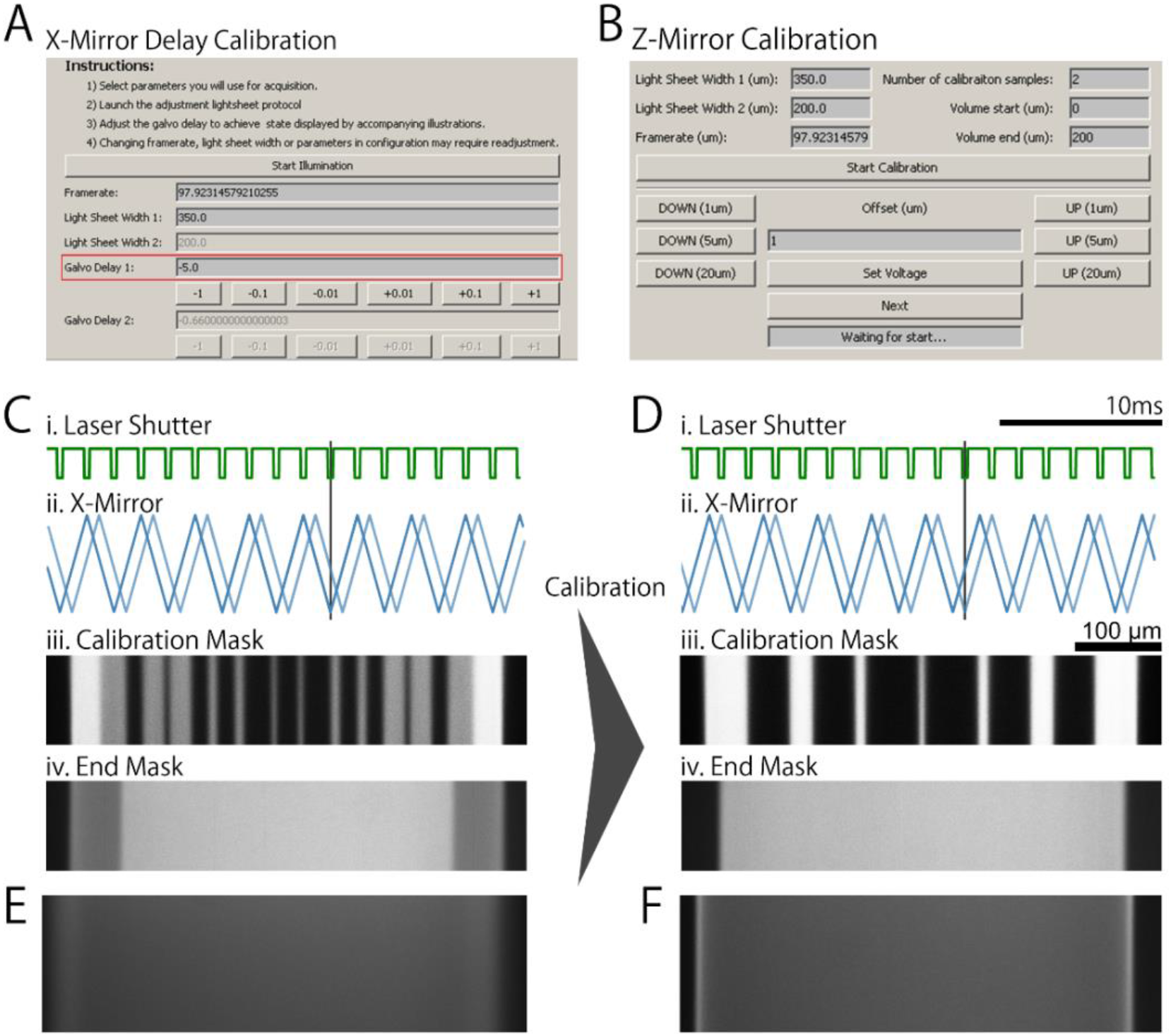
μSPIM Calibration. μSPIM provides the user with calibration tools which can be used to assess and correct the performance of the individual microscope elements. **(A)** and **(B)** show the calibration interface for X mirror with respect to the laser shutter when using masks and for Z mirror, respectively. **(C)** The movement of the X mirror galvanometer lags behind the supplied signal, resulting in artefacts when using laser masks. **(i.)** shows the laser shutter signal (green) and **(ii.)** shows the X mirror signal (blue) and actual mirror movement (light blue). Black bar highlights the misalignment between the shutter signal and mirror movement when the X mirror signal is not calibrated and no delay is introduced. **(iii.)** shows a mask used for calibration, emphasizing the difference between uncalibrated and calibrated system. **(iv.)** show the corresponding light sheet produced with poor calibration for the mask signal in (i.). **(D)** is analogous to **(C)**, showing a well calibrated system where the laser shutter **(i.)** is synchronized X mirror movement **(ii.)**, producing overlapping masks shown in **(iii.)** and **(iv). (E)** Uncalibrated Z-Mirror results in loss of focus through the volume. **(F)** After calibration, the light sheet is always in focus throughout the volume scan. All figures illustrating the light sheet in both uncalibrated and calibrated scenarios were acquired by scanning laser through a fluoroscein solution.

### A. X-Mirror

Proper X-plane mirror calibration is essential for the correct assessment of size of the illuminated region when starting acquisition. To calibrate the X mirror, it is necessary to know the size of the field of view (or pixel size). This can be calculated as the effective camera image sensor size divided by the magnification of the collection objective or by measuring a standard stage calibration slide under the microscope. Using this measurement, the X Mirror scale can be adjusted such that a sheet of a particular width matches the corresponding number of the pixel. In the case of two illumination paths, this procedure should be repeated for both paths separately. This calibration should be carried out such that the x-mirror moves around its central position when energized (i.e. with zero volts driving signal) and the illumination arm should be adjusted such that this central position is centred on the field of view.

### B. Piezoelectric Stage

Piezoelectric stages are generally factory-calibrated to move a predefined distance by set voltage difference (e.g. 1 μm per 40 mV for the PIFOC model) to which the Piezo Scale setting should be set. The unit is mV/μm and for the previous example this setting would be set to 40.0. μSPIM also provides a Piezo Offset setting which can be used to adjust voltage offset in software. For optimal operation, it is suggested to adjust the offset such that a plane at 0.0 μm is near the top of the available range. Setting the offset could be beneficial, for instance, when imaging at multiple depths in a sample so that when 0.0 μm is positioned at the top of the sample it serves as a reference while allowing the full range of the piezoelectric stage to be utilized.

### C. Z-Mirror

Next, it is necessary to calibrate the relative movement of the Z-plane mirror(s) with respect to the movement of the now-calibrated Piezoelectric stage. This can be done using a calibration procedure provided by μSPIM which steps the stage through a user-defined number of levels across a volume of choice and allows the user to adjusts the Z mirror voltage for each level such that the laser sheet is in focus. Finally, a Z Mirror scale is generated which can then be used to adjust settings.

### D. X-Mirror Lag Compensation

The X Mirror movement lags the signal which can be problematic when using laser masks (discussed in Section VI-A1). To correct for this, μSPIM provides a simple calibration process during which a distinct mask is displayed and the mirror lag is adjusted by the user until the masks perfectly overlap as shown in **Figure 7**.

**Fig. 7:**
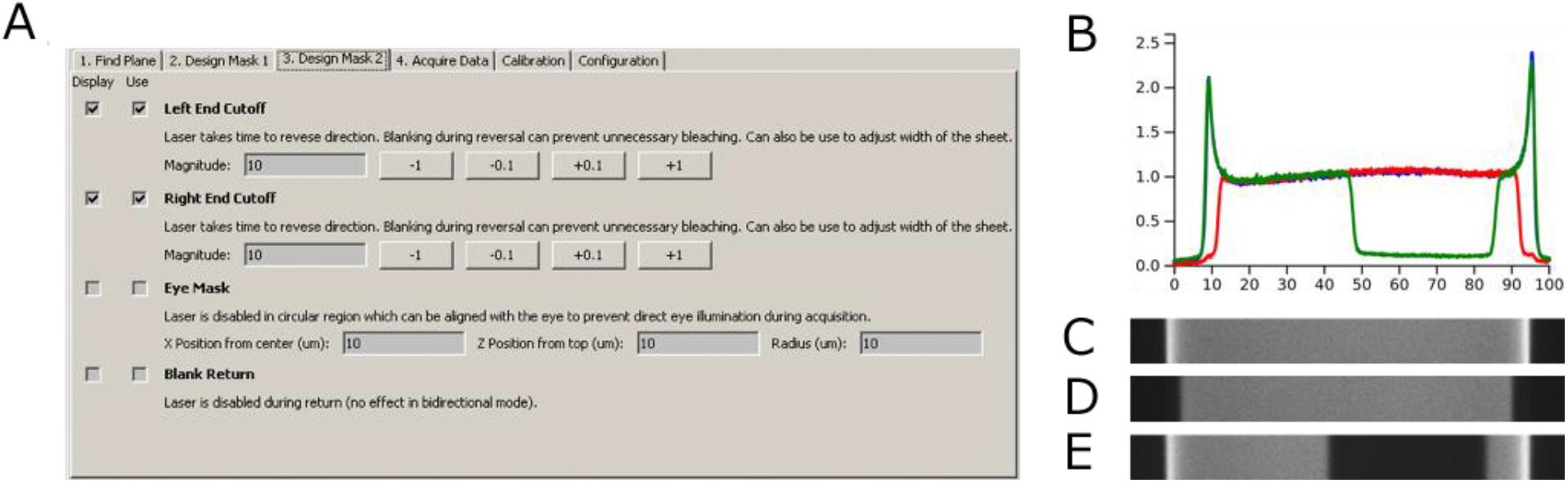
Light-Sheet Laser Masks. **A.** User Interface used to define laser masks allowing the user to display and use a number of masks for recordings: End cutoff Masks help eliminate excess light during laser return as visible in D (with cutoff masks enabled) compared to C (without any masks), Eye Mask allowing to turn the laser off in a specific region (such as eye of larval zebrafish) and Blank Return which turns the laser off during return (flyback), avoiding unnecessary photobleaching of the sample. **B** The median-normalized luminance intensity profiles of a light sheet in Fluorescein with no mask applied (blue, shown in C), edge cutoff masks (red, shown in D) and eye mask (green, shown in E).

## VI. Data Acquisition using μSPIM

For the acquisition of a single plane, the X mirror galvanometer has to be scanned at least once across the full range of the desired light sheet width and back to its original position (**Figure 2**, showing 3 scans per frame). To ensure uniform illumination between frames, μSPIM takes control over the camera using a rising edge trigger signal synchronized with the X mirror (**Figure 2**). Generation of a volume recording is provided by a synchronous movement of the Z Mirror and the Piezoelectric Stage between frames, acquiring consecutive frames from different heights in the samples (**Figure 2**).

μSPIM implements three volume acquisition modes: Scan Down, Scan Up and Bidirectional mode. In the Scan Down mode, the volume frames are acquired starting from the top, moving down through the volume and then finally returning to the top of the volume to prepare for the acquisition of the next volume. Since the stage moves a relatively heavy objective, instant movement from the bottom to the top of the volume is not possible and fast movement can introduce oscillations in the stage’s position, reducing the quality of the recording. To eliminate this issue, a flyback period is introduced at the end of volume acquisition (shown in **Figure 2B v**.), slowing down the return of the stage at the expense of several frames. The Scan Up mode offers similar functionality to that of Scan Down mode, however in the opposite direction. Bidirectional mode eliminates the need for the flyback period by alternating between Scan Down and Scan Up modes on consecutive volume acquisitions (**Figure 2B vi**.) at the expense of unequal time period between frame acquisitions.

To allow for better distribution of sample illumination or illumination of occluded regions, μSPIM also supports the control of two separate paths, allowing for two light sheets of different widths.

### A. Advanced Acquisition Methods

While basic acquisition described above may be sufficient for some applications, μSPIM is primarily intended for functional imaging with visible light which can disturb the live sample. To minimize the impact of the visible light, the μSPIM toolset implements several procedures which can be used in applications where this is an issue.

#### 1) Laser Masks

Removing parts of the light sheet can be beneficial in some acquisition settings. μSPIM utilizes fast software shuttering to provide two main masks which may be beneficial for *in vivo* imaging: *Edge Masks*, removing the ends of the light sheet and thus reducing the regions which are over-illuminated due to slow reversion of the direction of the X mirror (**Figure 7**) and *Eye Mask* which can be used to remove illumination from a selected region in the light sheet (**Figure 7**), allowing for acquisition where a specific region must not be illuminated (such as the eye).

#### 2) Light Adaptation

Zebrafish larvae have been shown to react to sudden light changes(Burgess and Granato, 2007), interfering with behaviours of interest. This proves problematic in visible light recordings as the light is commonly turned off between recordings to prevent photodamage to the sample, resulting in a sudden light change during the start of the recording. We provide a simple light adaptation functionality which places a light sheet in a position near the top of the sample prior to recording where photodamage is of less concern and then move the sheet into position as the recording is started.

#### 3) Multi-recording sequence

Similarly, to the adaptation period, sudden light changes can be reduced by consecutive acquisition of a number of recordings without the need for user input in between the recordings. This eliminates the concern of photodamage during non-acquisition time periods, allowing the laser to be turned on for the whole sequence of the recordings.

### B. Data Output Format

The image data output follows the MicroManager format with acquired image sequences being split into 4GB files accompanied by a metadata file (Metadata format can be found in the official MicroManager documentation: https://micro-manager.org/wiki/Files_and_Metadata). To accommodate for the extra information regarding the light sheet acquisition, μSPIM provides a separate metadata file which carries the information about the volume acquired as well as information about the experiment and sample provided by the user (**Figure 8**).

**Fig. 8:**
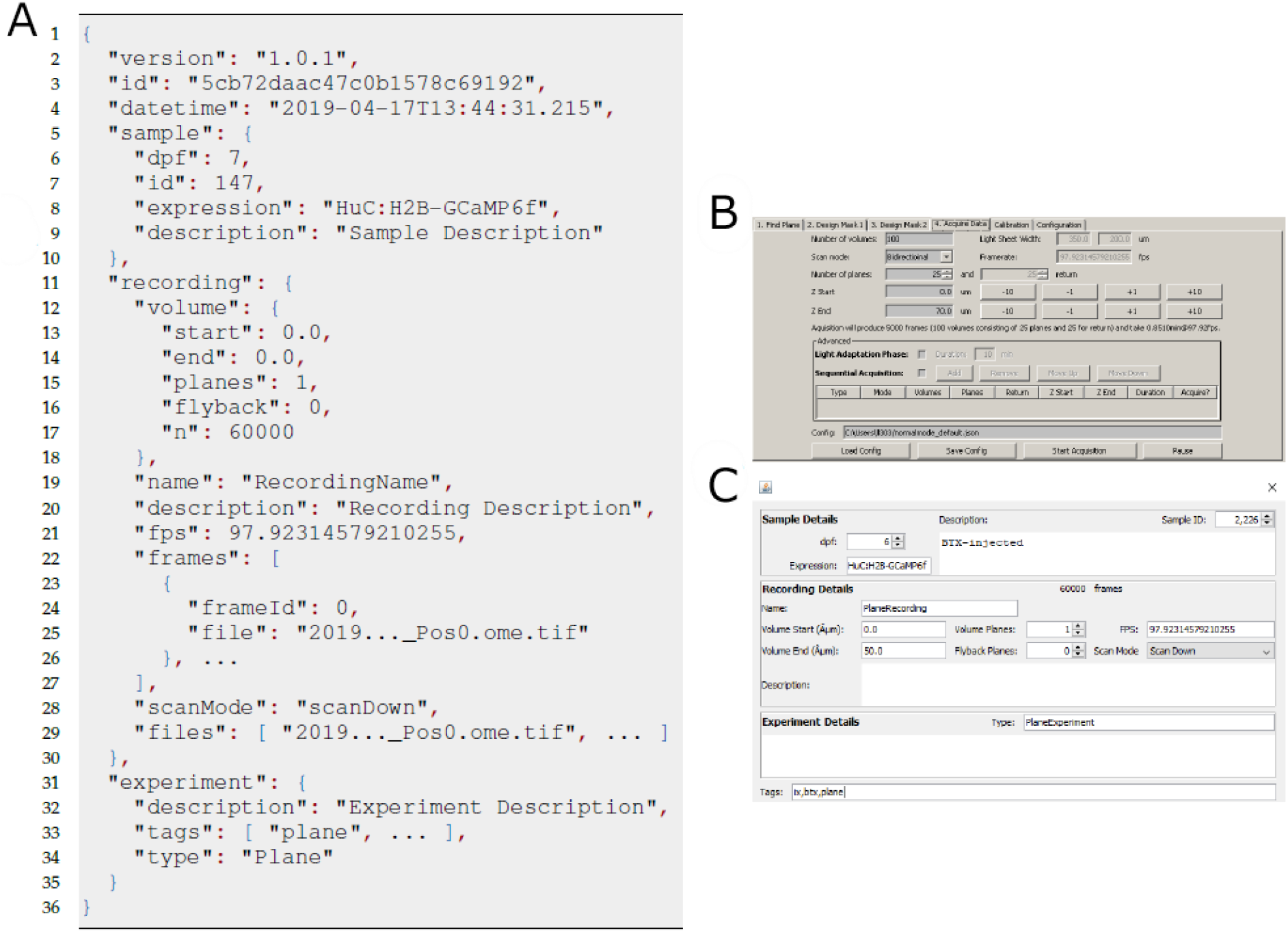
μSPIM Acquisition & Data Format. μSPIM generates a separate metadata file (shown in A), supplementing metadata supplied by the MicroManager platform. The generated data contains information detailing the settings of the acquisition (such as the height of the recorded plane or number of planes in the recorded volume) based on the parameters set by the user in B. This information can be supplemented by the user to contain information about the sample imaged, comments about the experiment protocol as well as the recording itself to aid cataloguing of the data.

## VII. Discussion

We have presented the μSPIM toolset which provides a flexible software solution for the control of SPIM microscopes and demonstrated its utility for two important applications – brain-wide imaging of neural activity in larval zebrafish (Ahrens et al., 2012) and visualizing neurons with cellular resolution in whole mouse brains (Tomer et al., 2014). In contrast to other open source control solutions for light sheet microscopes, such as OpenSPIM (Pitrone et al., 2013) and Open SPIM microscopy (Gualda et al., 2013), μSPIM focuses on a microscope implementation better suited for functional imaging (as opposed to developmental imaging), filling a gap in available software solutions rather than competing with the existing implementations. With a range of in-built calibration protocols, μSPIM allows the user to select hardware based on the needs of the given application. The MicroManager platform on which μSPIM is built provides support for a wide range of acquisition hardware which will provide users with a range of options in customizing their instrument. Comprehensive documentation and the open source nature of the toolbox allows the user to adapt the software to advanced application and non-standard use-cases, overcoming the current limitations of the software.

While our framework offers great flexibility in the choice of hardware used, and is modular allowing incomplete implementations, μSPIM fundamentally depends on the National Instruments DAC and other digital-analogue converters are currently not supported. Adding support for other DAC devices is supported by the structure of the source code of our control software, but the modifications necessary may be quite extensive.

As previously outlined, the focus of the μSPIM toolbox is primarily on the hardware flexibility aimed at a particular application (functional light sheet imaging) and thus some more complex functionality may not be supported out of the box. One notable example would be use of multiple lasers with various wavelengths which is required when imaging with multiple fluorescent reporters or markers: μSPIM is currently limited to providing support for two independent lasers. We have not yet explored the possibility of replacing one or either of these by a multi-colour light engine, but envisage that this will be possible given that micromanager supports a number of such devices. Again, advanced users may choose to modify our solution to support this functionality if required.

Being built around MicroManager, μSPIM inherits all of its limitations. For instance, while the list of acquisition hardware supported by the MicroManager platform is extensive, it is possible that some less common devices are not supported by the platform and thus cannot be used with our toolbox. Another notable limitation is the lack of support for the “rolling shutter” mode of acquisition in which the exposure of each line of pixels on the sCMOS sensor is delayed to coincide with the time at which those pixels are parfocal with the laser beam passing through the sample (also called electronic confocal slit detection (Hu et al., 2017)). Because only that single line of pixels is activated, much background fluorescence caused by scattered illumination of neighbouring areas is rejected, thereby improving resolution. We did not implement this mode of acquisition as a standard feature of μSPIM because it results in lower frame rates compared to the usual “global shutter” mode of acquisition in which all pixels are activated simultaneously.

While our framework does not support some use cases in its current implementation, it provides a comprehensive solution to the control of light sheet microscope with support for a wide range of both control and acquisition hardware while retaining a gentle learning curve supported by the well documented calibration and acquisition protocols. For this reason, we hope the μSPIM toolbox will be adopted in both standard and custom light sheet imaging use cases thanks to well-documented open source code base. Both documentation and source code is available for free from the μSPIM git repository (https://uspim.org).

## VIII. Materials and Methods

### A. Optical design of SPIM used in this work

A partial parts list for the SPIM we constructed is provided in Table 1. The illumination arm (Fig. 1) was fed by one or two lasers through optical fibres (kineflex). When using both red and green fluorophores, a long-pass dichroic mirror (Thorlabs DMLP505R) combined the beams of the blue and yellow lasers, which then passed through a mechanical shutter. The beam was reflected by a pair of X and Z Galvo scanning mirrors with 1 kHz bandwidth at deflection angles ± 0.2°. The beam was then expanded by a factor of 2.5 using a pair of achromatic lenses (f = 50 mm and 125 mm) The optimum beam parameters were calculated using calctool (http://www.calctool.org/CALC/phys/optics/f_NA). The illumination lens was f = 40 mm achromat. The whole of the illumination arm, beginning with the combining mirror, was mounted on an optical rail which was itself mounted on two translation stages (Thorlabs XR25C/M), one of which was slaved to the other. This arrangement allowed centering of the illumination beam on the field of view when the x-mirror was set to its position of zero offset, which is especially important for calibration of x-mirror displacement described above. With the rapid scanning of the X Galvo mirror, the laser line will form an illumination plane over the integration time of a single frame. The thickness of the beam waist was ∼7 μm, with a Rayleigh length of ∼440 μm. We therefore achieved a relatively uniform plane of illumination across half of the total field of view, which was 810 μm wide. The specimen stage was custom-designed according to application and manufactured using a 3D printer. The stage was attached to an x-y-z translation assembly (Thorlabs PT3) for positioning of the specimen.

**Table 1.** Parts list for construction of SPIM.

| **Mechanical Parts** | **Component (Manufacturer)** |
| --- | --- |
| Optical table | 1.4 x 1.8 x 0.2 m clean top on Micro-g Pneumatic and Rigid Legs (TMC Vibration Control) |
| Rail systems | Precision 100 mm Dovetail Optical Rails with PRC carriers (Newport) |
| Translation stages | XR25C/M (Thorlabs) |
| **Illumination arm** |  |
| Lasers | LuxX+ 488 nm 60 mw (Omicron)<br>Jive 561 nm 100 mW (Cobolt) |
| Optical Fibre and laser launch | Kineflex Fibre system (0.7 mm), 400-640nm, 2m, FC/APC connectors (Qioptiq) |
| Mechanical Shutter | SH05/M shutter and SC10 controller (Thorlabs) |
| Beam expander 2.5x | f = 50 mm achromat (AC254-050-A, Thorlabs) and f = 125 mm achromat (Edmund Optic) |
| Dual-axis galvanometer scanner | GVS102 - 2D Galvo System (ThorLabs) |
| Illumination lens | f = 40 mm achromat (#49-354-INK, Edmund Optics) |
| **Collection arm** |  |
| Camera | ORCA-flash4.0 v2 |
| Laser cleanup notch filters | 488/10, and 561/10 (Chroma) |
| Detection objective | 16X/0.8NA CFI LWD Plan Fluorite (Nikon) |
| Piezo | PIFOC E-665 amplifier/controller + P-725 (400 um travel range from Physike Instrumente) |
| Emission filter | 535/35 (xf3007, Omega) |
| Tube lens | f= 200 mm achromat (AC508-200, Thorlabs) |
| Mounting of collection arm | X-95 mounting system and carriers (Linos) |
| **Computing** |  |
| Custom built PC incorporating Intel® Xeon® processor E5-2600 |  |
| Frame Grabber | Firebird Camera Link (2xCLM-2PE8) (Active Silicon) |
| DAC card | PCIe-6738 (National Instruments) |

For detection, a 16X/0.8NA Nikon CFI LWD Plan Fluorite objective was placed perpendicularly to the illumination plane to collect the emitted fluorescence signal. The excitation light was rejected by the emission filter and then a tube lens of 200 mm focal length (AC508-200-A-ML, Thorlabs, Inc) used to project an image onto the sensor of the Hamamatsu Flash4.0 sCMOS camera (13 x 13 mm CCD, so that each pixel imaged an area of ∼0.16 μm2 using a 16x objective). The objective lens was mounted on a piezoelectric stage (P-721 PIFOC High-Precision Objective Scanner, Physik Instrumente Ltd) and its movement synchronized with the Z Galvo scanner to make sure the illumination plane is always in the imaging focal plane. The objective and PIFOC scanner were themselves mounted on a manual stage for finding the focal plane before starting acquisition.

### B. Imaging Brains of Larval Zebrafish

All procedures were in accordance with the UK Animal Act 1986 and were approved by the Home Office and the University of Sussex Ethical Review Committee. Transgenic zebrafish larvae of undetermined sex expressing the nuclear localised H2B-GCaMP6f calcium reporter panneuronally (Tg(elavl3:H2BGCaMP6f) (Chen et al., 2013; Dunn et al., 2016)), were imaged at 6 days-post-fertilisation (dpf). Larvae were paralysed by positioning them sideways in a small slit of PDMS (Sylgard184, Dow Crowning) on a coverslip (Pichler and Lagnado, 2019) and injecting 0.25mM a-Bungarotoxin (Tocris Bioscience) into their heart. They were then embedded dorsal side up in 2% low-melting-point agarose (Biogene) in E2 medium (Brand et al., 2002) a 22 mm × 22 mm coverslip and placed into a custom glass-walled 3D printed chamber. After filling the chamber with E2 medium, the majority of the agarose that fell between the laser source and the larva’s head was removed with a scalpel to maintain sufficient stability of the sample while minimising the way the laser travelling through the agarose. The chamber was positioned underneath the objective so that upon turning on the laser, it would not hit the head, and the eyes in particular, but rather the tail of the fish. Once imaging and excitation plane were successfully aligned, the larva’s brain was approached and the desired brain region was found, the size of the volume and the number of slices were chosen.

### C. Imaging Cleared Mouse Brains

Expression of the calcium-sensitive reporter GCaMP6f in pyramidal neurons of primary visual cortex was driven under the CaMKII promoter by injection of a virus (AAV1.CaMKII.GCaMP6f.WPRE.SV40 virus; Cat. No. 100384-AAV1, Addgene; titer 4.1011 GC/ml) to achieve a final concentration of 4mg/μl. titer 4.1011 GC/ml). Surgery was performed as described (Ryan et al., 2020). All experimental procedures were conducted according to the UK Animals Scientific Procedures Act (1986). Experiments were performed at University of Sussex under personal and project licenses granted by the Home Office following review by the Animal Welfare and Ethics Review Board. Tau preformed fibrils were prepared as previously described (Iba et al., 2013), (Peeraer et al., 2015) and kindly provided to us by Kristof Van Kolen (Janssen Pharmaceutica). Before injection these preformed fibrils were mixed with AAV virus to initiate tauopathy. All these procedures were carried out in mice of mice transgenic for the 383 aa isoform of human tau with the P301S mutation associated with frontotemporal dementia (Allen et al., 2002).

Mice were perfused transcardially with PBS, followed by freshly made 4% PFA in PBS. Brains were removed and kept in 4% PFA at 4°C overnight with shaking before washing in PBS (30min x 3). Brains were treated according to the iDISCO protocol (Renier et al., 2014). Briefly, the PFA fixed brains were gradually dehydrated with methanol in PBS (20%, 40%, 60%, 80%, 100%; 1 hour each) and then bleached in a 1:5 ratio of 30% hydrogen peroxide:methanol at 4°C overnight. Brains were rehydrated with methanol in PBS (80%, 60%, 40%, 20%, PBS; 1 hour each), and washed twice in PBS with 0.2% Triton X-100 (1-hour x 2, at room temperature). Brains were then treated in the permeabilization solution (PBS with 0.2% Triton X-100, 0.3M glycine, 20% DMSO) for 2 days with shaking at 37°C, followed by incubation in the blocking solution (PBS with 0.2% Triton X-100, 6% goat, 10% DMSO) for 4 days with shaking at 37°C. Antibody labelling and washes were performed in the same solution (PBS with 0.2% Tween-20 and 10ml/ml heparin). GFP was immunolabelled with an anti-GFP antibody (ab290, 1:200), and tau with the AT8 antibody (in-house, 1:200) (6-day-incubation with shaking at 37°C). The secondary incubation was performed using goat anti-rabbit (A11008, Invitrogen, 1:200, labelled with Alexa Fluor 488) and goat anti-mouse (A11004, Invitrogen, 1:200, labelled with Alexa Fluor 568) secondary antibodies (6-day-incubation with shaking at 37°C), each with addition of 3% goat serum. Excess primary and secondary antibodies were washed overnight with shaking at room temperature. Brains were dehydrated in methanol:H2O (20%, 40%, 60%, 80%, 100% x 2) and were left in methanol overnight. Lipids were extracted in 66% dichloromethane (DCM)/33% methanol at room temperature (3-hour incubation with shaking). Samples were washed in 100% DCM (15min x 2 with shaking). Brains were cleared using dibenzyl ether (DBE). Tubes were filled to the top to avoid oxidation and prevent brains turning opaque. Brains were kept in DBE-filled tubes until imaged. Red emission was collected through a 607/42 filter (Edmund Optics). Laser powers were usually ∼20 mW.

